# Decreased Left Atrial Cardiomyocyte FGF13 Expression Increases Vulnerability to Postoperative Atrial Fibrillation in Humans

**DOI:** 10.1101/2024.01.30.577790

**Authors:** Matthew A. Fischer, Adrian Arrieta, Marina Angelini, Elizabeth Soehalim, Douglas J. Chapski, Richard J. Shemin, Thomas M. Vondriska, Riccardo Olcese

## Abstract

Postoperative atrial fibrillation (POAF) is the most common complication after cardiac surgery and a significant cause of increased morbidity and mortality. The development of novel POAF therapeutics has been limited by an insufficient understanding of molecular mechanisms promoting atrial fibrillation. In this observational cohort study, we enrolled 28 patients without a history of atrial fibrillation that underwent mitral valve surgery for degenerative mitral regurgitation and obtained left atrial tissue samples along the standard atriotomy incision in proximity to the right pulmonary veins. We isolated cardiomyocytes and performed transcriptome analyses demonstrating 13 differentially expressed genes associated with new-onset POAF. Notably, decreased expression of fibroblast growth factor 13 (FGF13), a fibroblast growth factor homologous factor known to modulate voltage-gated sodium channel Na_V_1.5 inactivation, had the most significant association with POAF. To assess the functional significance of decreased FGF13 expression in atrial myocytes, we performed patch clamp experiments on neonatal rat atrial myocytes after siRNA-mediated FGF13 knockdown, demonstrating action potential prolongation. These critical findings indicate that decreased FGF13 expression promotes vulnerability to POAF.

---

Postoperative atrial fibrillation (POAF) complicates 30-40% of cardiac surgeries and increases short- and long-term morbidity and mortality^1^. Advancement in developing novel POAF therapeutics has been prevented by an inadequate understanding of molecular mechanisms promoting vulnerability to atrial fibrillation (AF)^1^. In this observational cohort study, we performed transcriptome analyses in left atrial myocytes from patients undergoing mitral valve surgery and identified differential gene expression associated with new onset POAF. Notably, the most significant association with POAF was decreased expression of fibroblast growth factor 13 (FGF13)^2^, the depletion of which we demonstrate is sufficient to prolong the action potential in rat atrial myocytes.

After IRB approval and informed consent, we enrolled adult patients without a history of AF presenting for elective mitral valve surgery for severe degenerative mitral regurgitation. Patients with prior cardiac surgery, mitral stenosis, aortic regurgitation, aortic stenosis, and endocarditis were excluded. Patients were consecutively enrolled at Ronald Reagan UCLA Medical Center between April 2021 and June 2022. POAF was defined as any clinician diagnosis of AF by electrocardiogram of at least 30s duration from ICU admission until hospital discharge. All patients had continuous electrocardiogram monitoring throughout their hospital course, which was independently reviewed daily for validation of POAF diagnosis. Patients did not receive prophylactic amiodarone. Among 28 patients enrolled with no history of AF (**Figure 1A**), 15 experienced POAF (53.6%), an incidence consistent with the increased risk of POAF after mitral valve surgery^1^. Preoperative QTc prolongation was the only clinical characteristic significantly associated with POAF (p-value=0.04, t-test). Separately, we enrolled 2 patients with preexisting persistent AF presenting for the same surgery for qualitative comparison of POAF-associated gene expression.

**Figure 1.**
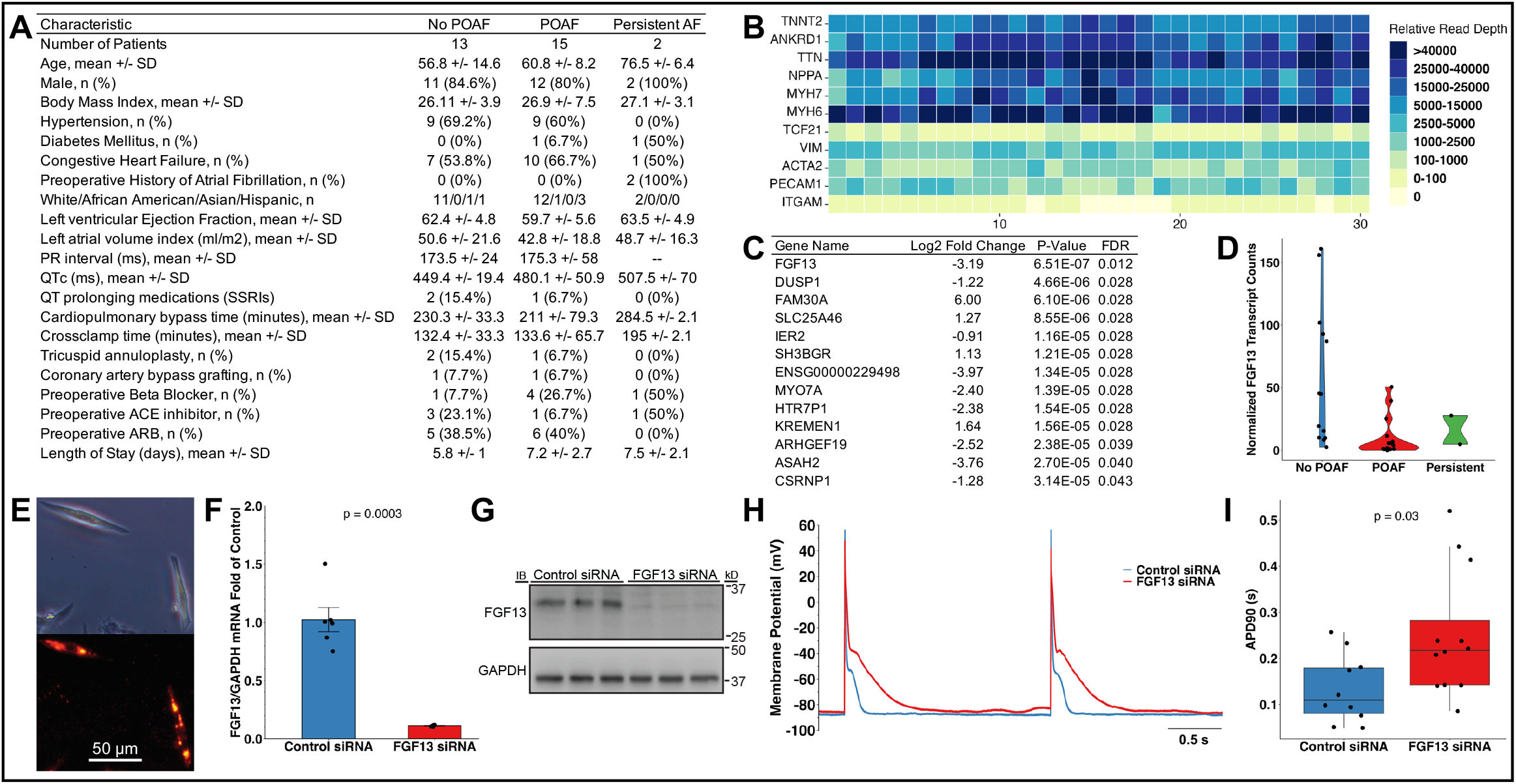
Decreased FGF13 mRNA expression in human left atrial cardiomyocytes promotes new onset postoperative atrial fibrillation. **A)** Clinical characteristics of enrolled patients. Preoperative QTc prolongation was the only clinical characteristic significantly associated with POAF (p=0.04, t-test). **B)** Heatmap of RNA-seq data shows enrichment for cardiomyocyte specific genes. Columns represent patient samples while rows correspond to individual genes. The top 6 rows correspond to 5 cardiomyocyte specific genes (TTNT2, ANKRD1, TTN, MYH7, MYH6) and an atrial specific gene (NPPA) while the bottom 5 rows correspond to fibroblast (TCF21, VIM and ACTA2), endothelial cell (PECAM1) and macrophage (ITGAM1) specific genes. **C)** Gene expression was compared between patients that experienced new onset POAF after surgery and those that did not. Genes with a statistically significant association with POAF (FDR < 0.05) are listed. Decreased expression of Fibroblast growth Factor 13 (FGF13), known to modulate voltage-gated sodium channel Na_V_1.5 inactivation, was the most significant association with POAF. **D)** Violin plots show normalized FGF13 transcript read counts in human left atrial cardiomyocytes, with decreased FGF13 expression associated with POAF. **E)** Brightfield microscopy and Cy3 fluorescence demonstrating transfection of FGF13 siRNA in neonatal rat atrial myocytes after 24 hr in culture **F)** RT-qPCR demonstrating 90% siRNA-mediated FGF13 knockdown 48 hours after transfection compared to negative control siRNA transfected myocytes. **G)** Immunoblot confirming FGF13 knockdown at the protein level 48 hours after transfection. **H)** Representative action potential recordings from control siRNA and FGF13 siRNA transfected atrial myocytes recorded at PCL 2s under current clamp configuration at room temperature. **I)** Current clamp experiments performed 48 hours after transfection show prolonged atrial myocyte action potential duration (p=0.03, Wilcoxon rank-sum test) after FGF13 siRNA-mediated knockdown (median APD_90_=0.217s, n=12) compared to control (median APD_90_=0.110s, n=10).

During cardiopulmonary bypass, the surgeon obtained a small (∼0.1g) sample of left atrial tissue along the standard incision on the left atrium at Waterston’s groove for surgical exposure of the mitral valve. Cardiomyocytes were isolated from frozen human left atrial tissue samples as previously described^3^. RNA was extracted from cardiomyocytes using the Qiagen AllPrep Micro Kit. RNA libraries were sequenced on an Illumina NovaSeq S2 instrument. Read pairs were pseudoaligned to hg38 using Salmon. We used DESeq2 to identify differential gene expression in our transcriptome-wide analysis (FDR<0.05, Benjamini-Hochberg procedure). RNA-seq analyses of cardiomyocytes demonstrated enrichment in markers of this cell type (**Figure 1B**), indicative of sample enrichment. Differential gene expression analyses revealed 13 genes associated (FDR<0.05) with POAF (**Figure 1C**). Among these, FGF13 was most significant (**Figure 1D**). The association of decreased FGF13 expression with POAF was highly significant (p-value=1.52E-08, FDR=0.0001) after controlling for age, sex and QTc. Fibroblast growth factor 13 (FGF13), a fibroblast growth factor homologous factor expressed in cardiomyocytes, is known to modulate voltage-gated sodium channel Na_V_1.5 inactivation^2^. The two patients with preexisting persistent AF had FGF13 expression levels comparable to the POAF patients. Given our POAF incidence, we were adequately powered (80%) to detect effect sizes ≥1.1 between groups (alpha=0.05, two-tailed) using the two-sample t-test as a simplification of the DESeq2 negative binomial model.

To isolate neonatal rat atrial myocytes, atria were dissected from euthanized P3 Sprague-Dawley rats, minced, and enzymatically digested. Atrial myocytes were isolated using a Percoll gradient and plated on 24-well plates containing 5mm coverslips coated with fibronectin. 24hrs later, atrial myocytes were co-transfected with Cy3-conjugated non-targeting siRNA and either FGF13-targeted or negative control siRNA for 12hrs. 48hrs later, transfected atrial myocytes were identified by Cy3 fluorescence followed by whole-cell patch clamp experiments to measure APD at 90% repolarization (APD_90_) at 2s cycle length or RT-qPCR/immunoblot to confirm FGF13 knockdown. Current clamp experiments demonstrated that FGF13 knockdown (**Figures 1E, 1F** and **1G**) caused significant atrial myocyte APD prolongation (control siRNA median APD_90_=0.110s, interquartile range=0.081s-0.179s, n=10; FGF13 siRNA median APD_90_=0.217s, interquartile range=0.142s-0.282s, n=12; p=0.03, Wilcoxon rank-sum test; **Figures 1H and 1I**).

FGF13 has been shown in animal models to be protective in inhibiting pathological late sodium current^4^, which has been implicated in AF pathogenesis by prolonging APD. Accordingly, FGF13 knockout was also shown to increase the late sodium current^5^. In this study, we provide critical insight into the relevance of FGF13 dysregulation to acquired arrhythmia in humans: 1) we measure FGF13 mRNA in human left atrial samples obtained in proximity to the pulmonary veins, the most common anatomical origin of AF; 2) we link decreased FGF13 levels to new onset POAF; 3) we show that FGF13 expression is decreased in POAF patients *prior to the onset of AF*, providing a clinically actionable target for POAF prophylaxis; and 4) we demonstrate APD prolongation in atrial myocytes after FGF13 knockdown, validating a molecular mechanism for arrhythmia. Our findings support a model whereby decreased FGF13 expression creates vulnerability to AF through left atrial myocyte APD prolongation that, following the stress response to surgery, precipitates AF in the postoperative period. These critical findings support further research in the use of selective late sodium current inhibitors for POAF prophylaxis.

## Acknowledgements

We thank the UCLA Cardiac OR team for their support of this research, Tristan Grogan for performing power calculations and the UCLA Technology Center for Genomics & Bioinformatics for nucleotide sequencing.

